# Branched-Chain Amino Acid Metabolism is Regulated by ERRα in Primary Human Myotubes and is Further Impaired by Glucose Loading in Type 2 Diabetes

**DOI:** 10.1101/2020.07.24.218099

**Authors:** Rasmus J.O. Sjögren, David Rizo-Roca, Alexander V. Chibalin, Elin Chorell, Regula Furrer, Shintaro Katayama, Jun Harada, Håkan K.R. Karlsson, Christoph Handschin, Thomas Moritz, Anna Krook, Erik Näslund, Juleen R. Zierath

## Abstract

**Aims/hypothesis:** Increased levels of branched-chain amino acids (BCAAs) are associated with type 2 diabetes pathogenesis. However, most metabolomic studies are limited to an analysis of plasma metabolites under fasting conditions, rather than the dynamic shift in response to a metabolic challenge. Moreover, metabolomic profiles of peripheral tissues involved in glucose homeostasis are scarce and the transcriptomic regulation of genes involved in BCAA catabolism is partially unknown. This study aimed to identify differences in circulating and skeletal muscle BCAA levels in response to an OGTT in individuals with normal glucose tolerance (NGT) or type 2 diabetes. Additionally, transcription factors involved in the regulation of the BCAA gene set were identified.

**Methods:** Plasma and *vastus lateralis* muscle biopsies were obtained from individuals with NGT or type 2 diabetes before and after an OGTT. Plasma and *quadriceps* muscles were harvested from skeletal muscle-specific PGC-1α knockout and transgenic mice. BCAA-related metabolites and genes were assessed by LC-MS/MS and RT-PCR, respectively. Small interfering RNA and adenovirus-mediated overexpression techniques were used in primary human skeletal muscle cells to study the role of *PGC-1A* and *ESRRA* in the expression of the BCAA gene set. Radiolabeled leucine was used to analyze the impact of ERRα knockdown on leucine oxidation.

**Results:** Impairments in BCAA catabolism in people with type 2 diabetes under fasting conditions were exacerbated after a glucose load. Branched-chain keto acids were reduced 37–56% after an OGTT in the NGT group, whereas no changes were detected in individuals with T2D. These changes were concomitant with a stronger correlation with glucose homeostasis biomarkers and downregulated expression of BCAT2, BCKDH complex subunits and 69% of downstream BCAA-related genes in skeletal muscle. In primary human myotubes overexpressing PGC-1α, 61% of the analyzed BCAA genes were upregulated, while 67% were downregulated in the *quadriceps* of skeletal muscle-specific PGC-1α knockout mice. *ESRRA* (encoding estrogen-related receptor α, ERRα) silencing completely abrogated the PGC-1α-induced upregulation of BCAA-related genes in primary human myotubes.

**Conclusions/interpretation:** Metabolic inflexibility in type 2 diabetes impacts BCAA homeostasis and attenuates the decrease of circulating and skeletal muscle BCAA-related metabolites after a glucose challenge. Transcriptional regulation of BCAA genes in primary human myotubes via a PGC-1α is ERRα-dependent.

**Research in context:** *What is already known about this subject?:*

- Circulating levels of BCAA are elevated in type 2 diabetes.
- PGC-1α is involved in the transcription of the BCAA gene set.

*What is the key question?:*

- Does metabolic inflexibility associated with type 2 diabetes encompass BCAA homeostasis and PGC-1α mediated transcription of the BCAA gene set?

*What are the new findings?:*

- BCAA homeostasis is further compromised by a glucose challenge in type 2 diabetes.
- An OGTT reveals coordinated regulation between BCAA metabolites, blood glucose, and HbA_1c_ levels.
- ERRα is essential for PGC-1α-mediated BCAA gene expression in primary human myotubes.

*How might this impact on clinical practice in the foreseeable future?:*

- An OGTT can be used to underscore impairments in BCAA metabolism. These findings suggest that interventions targeting the PGC-1α/ ERRα axis may improve BCAA homeostasis.

## Introduction

Type 2 diabetes is a metabolic disease characterized by chronic hyperglycemia and insulin resistance [1]. These metabolic derangements severely affect pathways controlling the appropriate sensing and handling of nutrients, thereby leading to a positive energy balance and metabolic inflexibility, which further compromises whole body glucose homeostasis [2]. Overnutrition and type 2 diabetes-related disturbances also affect non-glucose metabolites such as branched-chain amino acids (BCAAs; leucine, isoleucine and valine) [3], essential amino acids whose utilization and metabolism are exquisitely regulated in healthy individuals. While BCAA and related metabolites play a role in protein synthesis, they also modulate liver gluconeogenesis and lipogenesis rates, cell growth, and nutrient signaling, and can enter the tricarboxylic acid cycle to produce energy [4]. Under pathological conditions, BCAAs, especially in a context of overnutrition, disrupt insulin sensitivity and secretion [5].

Circulating levels of BCAAs are elevated in individuals with obesity, insulin resistance and/or type 2 diabetes [6]. Metabolomic profiling of blood metabolites has revealed a signature of altered BCAA catabolism in obese individuals, with a strong association with insulin resistance [7]. BCAA-related metabolites are predictive of type 2 diabetes pathogenesis [8], as well as prognostic markers for intervention outcomes [9, 10]. Mendelian randomization analysis suggested a causal link between genetic variants associated to impaired BCAA catabolism and higher risk of type 2 diabetes [11]. Therefore, there is growing evidence that high levels of BCAAs and related intermediate metabolites are not only type 2 diabetes biomarkers, but also pathophysiological factors. However, many of these studies were conducted in individuals after an overnight fast, under conditions in which protein degradation in skeletal muscle, and a concomitant release of amino acids to support gluconeogenesis [12], could mask deeper alterations in BCAA homeostasis. Moreover, they do not provide insight into the dynamic shift in metabolism that occurs in response to nutritional challenges. An analysis of metabolomic signatures in fasted individuals with glucose tolerance or type 2 diabetes before and after a glucose challenge may provide insight into dynamic changes in BCAAs and related metabolites and the consequences of insulin resistance.

Skeletal muscle is the largest contributor to systemic BCAA oxidation [13] and therefore impairments in glucose and BCAA metabolism in myocytes have an impact on whole body metabolic homeostasis. While BCAA-related gene expression and oxidation rates are reduced in *vastus lateralis* muscle from individuals with insulin resistance [14], studies of BCAA metabolism in human skeletal muscle are scarce. BCAA catabolism occurs in the mitochondrial matrix, implicating alterations in mitochondrial proteins may influence BCAA metabolism. In transgenic mice overexpressing the mitochondrial biogenesis inducer peroxisome proliferator-activated receptor γ (PPAR γ) coactivator 1α (PGC-1α), BCAA levels are reduced in *gastrocnemius* muscle [15]. Conversely, administration of the PPARγ agonist thiazolidinedione rosiglitazone improves glycemic control and increases circulating levels of BCAAs in individuals with type 2 diabetes [16]. Nevertheless, mechanisms underlying the direct role of PGC-1α in the regulation of BCAA catabolism are unclear.

The aim of this study was to assess whether an OGTT can further reveal an impairment in BCAA metabolism in individuals with type 2 diabetes. We also aimed at identifying a PGC-1α transcriptional partner in the regulation of the BCAA gene program by using cell and mouse models with altered PGC-1α expression.

## Research Design and methods

### Subjects

Thirty-two men with normal glucose tolerance (NGT) and twenty-nine men with type 2 diabetes aged 44-69 years were recruited to participate in this study. All participants gave their informed consent. The study was approved by the regional ethical review board in Stockholm. The experimental procedures were conducted according to the Declaration of Helsinki. Participants underwent a clinical health screening consisting of clinical chemistry and anthropometric measurements (Table 1 and 2). Five individuals in the control group that exhibited impaired glucose tolerance and one individual diagnosed with type 2 diabetes that exhibited NGT were excluded from the study Fig. 1. Individuals with type 2 diabetes had higher blood glucose, Hb1Ac, and HOMA-IR, as well as higher BMI and waist circumference (Table 1 and 2). Individuals with type 2 diabetes included in the transcriptomic analysis were older than the normal glucose tolerant controls (Table 2). Individuals with type 2 diabetes were treated with metformin (n=23; daily dose range 500-3000 mg) and/or sulfonylurea (n=7; daily dose range 2-5 mg). Anti-diabetic medication was taken by the individuals with type 2 diabetes after the skeletal muscle biopsy collection.

**Figure 1.**
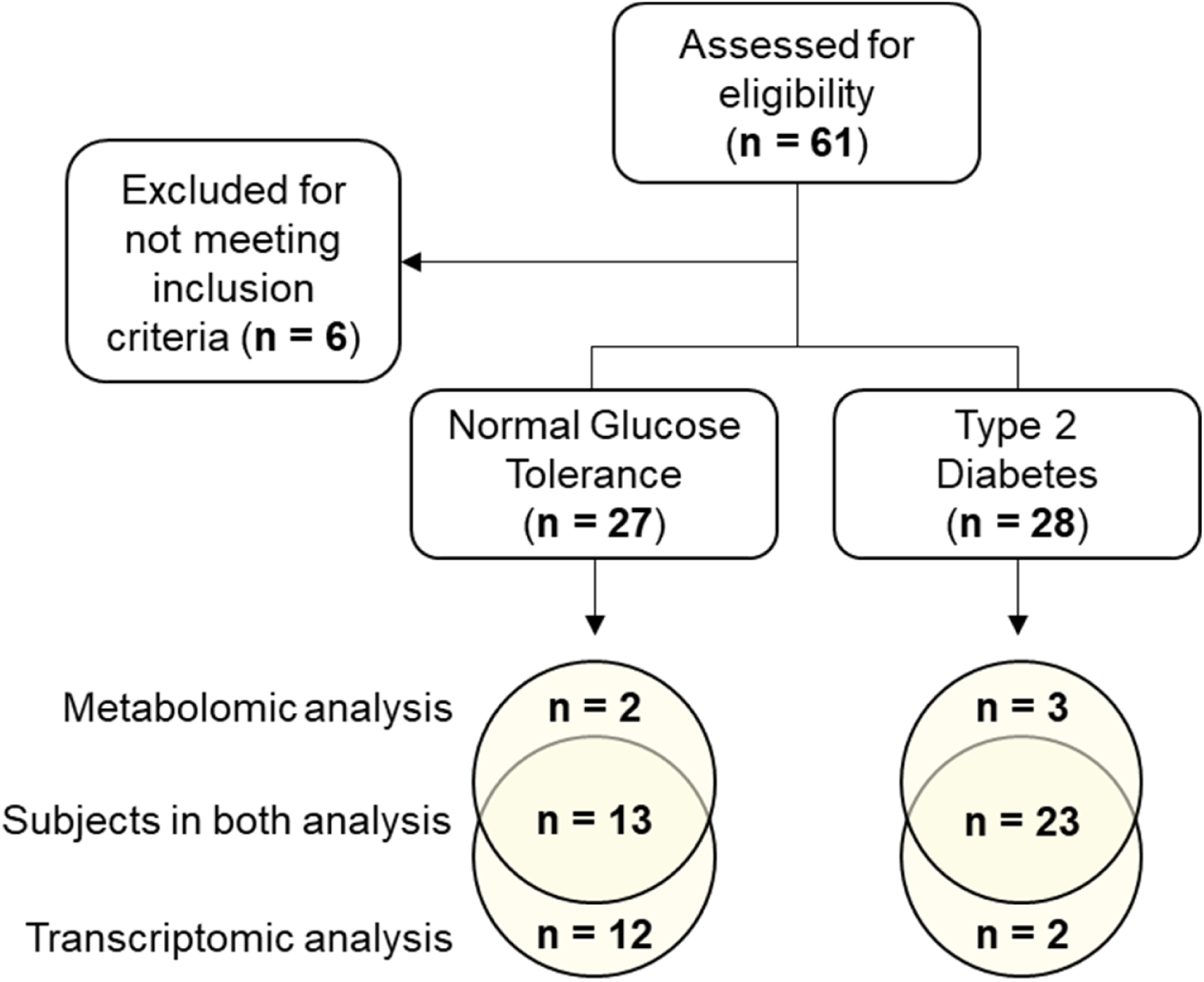
Flow chart illustrates subject enrolment and analysis. Sixty-one individuals were assessed for eligibility, six of which were excluded for not meeting the inclusion criteria. Group size for transcriptomic and metabolomic analysis is reported. Transcriptomic and metabolomic analysis shared samples from 13 individuals with normal glucose tolerance and 23 with type 2 diabetes subjects.

**Table 1.** Clinical characteristics of the subjects included in the metabolomic analysis.

|  | **Normal Glucose**<br>**Tolerance** | **Type 2**<br>**Diabetes** | **P-value** |
| --- | --- | --- | --- |
| **Age (years)** | 59 ± 2.4 | 62 ± 1.2 | 0.1611 |
| **Weight (kg)** | 81 ± 2.7 | 88 ± 1.8 | 0.0209 |
| **Height (m)** | 1.77 ± 0.02 | 1.79 ± 0.01 | 0.4082 |
| **BMI (kg/m<sup>2</sup>)** | 25.95 ± 0.59 | 27.66 ± 0.47 | 0.0307 |
| **Waist (cm)** | 90 ± 1.6 | 100 ± 1.6 | 0.0002 |
| **Hip (cm)** | 97 ± 1.2 | 101 ± 1.2 | 0.0241 |
| **WHR ratio** | 0.93 ± 0.01 | 0.99 ± 0.01 | 0.0008 |
| **Pulse (BPM)** | 62 ± 2.0 | 68 ± 1.6 | 0.0234 |
| **FP-glucose (mmol/L)** | 5.41 ± 0.13 | 8.59 ± 0.37 | <0.0001 |
| **120 min-glucose (mmol/L)** | 6.05 ± 0.25 | 16.66 ± 0.58 | <0.0001 |
| **HbA<sub>1c</sub> (mmol/mol)** | 36.13 ± 1.13 | 51.19 ± 1.32 | <0.0001 |
| **HbA<sub>1c</sub> (%)** | 5.5 ± 0.10 | 6.8 ± 0.12 | <0.0001 |
| **S-insulin (pmol/L)** | 52.37 ± 5.30 | 85.51 ± 9.06 | 0.0124 |
| **120 min S-Insulin (pmol/L)** | 294.2 ± 43.41 | 327.5 ± 42.06 | 0.6155 |
| **P-Creatinine (μmol/L)** | 84.53 ± 2.65 | 80.72 ± 2.17 | 0.2794 |
| **P-ASAT (μkat/L)** | 0.42 ± 0.04 | 0.39 ± 0.03 | 0.5992 |
| **P-ALAT (μkat/L)** | 0.38 ± 0.06 | 0.48 ± 0.04 | 0.1161 |
| **P-TG (mmol/L)** | 1.08 ± 0.14 | 1.41 ± 0.13 | 0.1093 |
| **P-Chol (mmol/L)** | 5.27 ± 0.17 | 4.50 ± 0.14 | 0.0014 |
| **P-HDL (mmol/L)** | 1.33 ± 0.06 | 1.24 ± 0.06 | 0.3562 |
| **P-LDL (mmol/L)** | 3.47 ± 0.13 | 2.62 ± 0.14 | 0.0003 |
| **S-C-peptide (nmol/L)** | 0.69 ± 0.04 | 0.99 ± 0.08 | 0.0115 |
| **HOMA1-IR** | 1.71 ± 0.22 | 5.28 ± 0.65 | 0.0013 |

Results are mean ± SEM for individuals with Normal Glucose Tolerance (n=15) and Type 2 Diabetes (n=26) men. Statistical analysis was performed using Student's *t*-test.

**Table 2.** Clinical characteristics of the subjects included in the transcriptomic analysis.

|  | **Normal Glucose**<br>**Tolerance** | **Type 2**<br>**Diabetes** | **P-value** |
| --- | --- | --- | --- |
| **Age (years)** | 58 ± 1.8 | 62 ± 1.3 | 0.0185 |
| **Weight (kg)** | 84 ± 1.7 | 89 ± 1.9 | 0.0314 |
| **Height (m)** | 1.80 ± 0.02 | 1.79 ± 0.01 | 0.7185 |
| **BMI (kg/m<sup>2</sup>)** | 25.98 ± 0.35 | 27.94 ± 0.48 | 0.0020 |
| **Waist (cm)** | 93 ± 1.4 | 101 ± 1.7 | 0.0020 |
| **Hip (cm)** | 99 ± 0.9 | 101 ± 1.3 | 0.1691 |
| **WHR ratio** | 0.94 ± 0.01 | 0.99 ± 0.01 | 0.0009 |
| **Pulse (BPM)** | 61 ± 1.4 | 68 ± 1.6 | 0.0072 |
| **FP-glucose (mmol/L)** | 5.46 ± 0.08 | 8.56 ± 0.39 | <0.0001 |
| **120 min-glucose (mmol/L)** | 5.89 ± 0.27 | 16.20 ± 0.69 | <0.0001 |
| **HbA<sub>1c</sub> (mmol/mol)** | 35.76 ± 0.64 | 50.44 ± 1.50 | <0.0001 |
| **HbA<sub>1c</sub> (%)** | 5.4 ± 0.06 | 6.8 ± 0.14 | <0.0001 |
| **S-insulin (pmol/L)** | 52.71 ± 5.58 | 92.58 ± 9.01 | 0.0008 |
| **120 min S-Insulin (pmol/L)** | 314.9 ± 50.8 | 350.5 ± 42.4 | 0.5937 |
| **P-Creatinine (µmol/L)** | 83.64 ± 2.36 | 79.71 ± 2.29 | 0.2307 |
| **P-ASAT (µkat/L)** | 0.38 ± 0.03 | 0.40 ± 0.03 | 0.7613 |
| **P-ALAT (µkat/L)** | 0.42 ± 0.05 | 0.50 ± 0.04 | 0.2565 |
| **P-TG (mmol/L)** | 1.25 ± 0.15 | 1.43 ± 0.13 | 0.4415 |
| **P-Chol (mmol/L)** | 5.40 ± 0.20 | 4.48 ± 0.15 | 0.0005 |
| **P-HDL (mmol/L)** | 1.30 ± 0.05 | 1.19 ± 0.06 | 0.1709 |
| **P-LDL (mmol/L)** | 3.56 ± 0.17 | 2.63 ± 0.15 | 0.0002 |
| **S-C-peptide (nmol/L)** | 0.69 ± 0.04 | 1.03 ± 0.08 | 0.0008 |
| **HOMA1-IR** | 1.86 ± 0.21 | 5.23 ± 0.64 | <0.0001 |

Results are shown as mean ± SEM for individuals with Normal Glucose Tolerance (n=25) and Type 2 Diabetes (n=25) men. Statistical analysis was performed using Student's *t*-test.

Participants arrived at the clinic at 07:45 after a 12-h fast. A catheter was inserted into an arm vein and fasting blood samples were obtained. After applying local anesthesia (Lidokain hydrochloride 5 mg/mL), biopsies from the *vastus lateralis* were obtained using a conchotome (AgnTho’s AB, Sweden). After ∼15 min, participants were given a solution containing 75 g of glucose and underwent an OGTT. A blood sample was obtained 30 min after glucose ingestion. After 120 minutes, a blood sample and *vastus lateralis* biopsy was obtained. All biopsies were s immediately snap-frozen in liquid nitrogen and stored at −80°C. Metabolomic analysis was performed by Metabolon, Inc. (Durham, NC) as described [17]. See electronic supplementary material [ESM] Methods: Human plasma and skeletal muscle metabolomics for details.

### Cells culture experiments

Human satellite cells were harvested from *vastus lateralis* skeletal muscle biopsies of healthy volunteers from both sexes and differentiated as described [18]. Mouse C2C12 myoblasts (ATCC CRL-1772, VA) were propagated in growth media (DMEM, 20% FBS, and 1% anti-anti). After ∼12 hours, medium was changed to differentiation medium (DMEM, 2% horse serum, and 1% anti-anti) and differentiated myotubes were cultured for six days before the final experiments. For pharmacological inhibition of ERRα, human skeletal muscle cells (HSMCs) were incubated with 5 μM of XCT-790 (X4753, Sigma-Aldrich), an inverse ERRα agonist, for 32 hours. Cells were regularly tested to confirm the absence for mycoplasma contamination.

Small interfering RNAs (siRNA) targeting *PPARGC1A* or *ESRRA* mRNA predesigned by ThermoFisher Scientific (Silencer® Select, s21395 and s4830) were used to silence the expression of these genes. Scramble siRNA (4390847, ThermoFisher) was used as negative control. Transfection of C2C12 myoblasts and HSMCs was performed 3 and 6 days after induction of differentiation, respectively, in OptiMEM reduced serum media (31985062, ThermoFisher) with Lipofectamine® RNAiMAX (13778, ThermoFisher).

Adenoviral-delivery of human and mouse PGC1α (Ad-PGC1α) or GFP (Ad-GFP) was performed over-night in differentiated HSMCs and C2C12 myotubes. Experiments were performed 48 hours after inducing transduction. In cells that were both treated with siRNA and transduced, silencing was performed as described above and cells were transduced immediately after transfection.

### Mouse models

Male mice were housed under controlled lighting (12 h light/12 h dark cycle) with free access to food and water. Experiments were performed in accordance with Swiss federal guidelines and were approved by the Kantonales Veterinäramt Basel-Stadt. Skeletal muscle-specific PGC-1α transgenic mice (mTG) are described elsewhere [19]. The PGC-1α muscle-specific knockout mice (mKO) were generated by breeding PGC-1α^loxP/loxP^ mice [20] with transgenic mice expressing the Cre recombinase under the control of the human α-skeletal actin promoter (HSA-Cre). Chow diet (AIN-93G; 7% fat, 58.5% carbohydrates, 18% protein and 16.5% of crude fiber, ash, and moisture) was provided by Provimi Kliba AG (Kaiseraugst, Switzerland).

Fasted (4h) male mice aged 11-14 weeks were anesthetized by intraperitoneal injection of pentobarbital (150 mg/kg) and tissues were collected. Metabolomic analysis of plasma and quadriceps muscle was performed at the Swedish Metabolomics Centre (Umeå University) as described [21]. See ESM Methods: Mouse plasma and skeletal muscle metabolomics for details.

### Real-time qPCR

Total RNA from human skeletal muscle biopsies, mouse skeletal muscles and cultured myotubes was extracted for mRNA analysis of BCAA genes. See ESM Methods: RNA isolation and relative mRNA expression for details. Primer sequences are shown in ESM Table 1 and ESM Table 2.

### Western blot analysis

Equal amounts of protein from human skeletal muscle biopsies (10 µg) and cells (20 µg) were analyzed for BCKDHA, p^S293^-BCKDHA, BCKDK and PPM1K. See ESM Methods: Western Blot analysis for details.

### Leucine oxidation

Myotube cultures were incubated with 1 ml of Ham’s F10 Medium in the presence of 0.1 mCi/ml radiolabeled leucine (L-[U-C^14^]-Leucine; 11.1GBq/mmol, NEC279E250UC, PerkinElmer) and either 3,6-dichlorobenzo(b)thiophene-2carboxylic acid (BT2) (ENA018104907, Sigma-Aldrich, Sweden) or DMSO. Small cups were placed in cell-culture wells and plates were air-tight sealed. After a 4-hour incubation at 37°C, 150 μL 2 M HCl and 300 μL of 2 M NaOH were added into each well and small cup, respectively. After 1 hour, the liberated CO_2_ was collected and subjected to scintillation counting (Tri-Carb 4910TR, PerkinElmer, MA, USA).

### Statistical Analysis

Results in tables are reported as means ± SEM. Metabolites and gene expression results are reported as Tukey boxplots with median (line), 25–75% (box) and the 25th/75th percentile minus/plus 1.5 times the inter-quartile distance (whiskers). Values outside this range are plotted as individual open circles. Outlier values detected using the Grubb’s test are plotted as closed circles and were not considered in the statistical analysis. HSMCs from each donor, C2C12 myotubes at different passages, and single animals, were considered as experimental units. Data normality was tested using the Shapiro-Wilk test. Homogeneity of variance was tested using the Levene’s test. Data that did not meet these criteria was transformed with Tukey’s Ladder before the significance testing. Data was analyzed using GraphPad Prism software (GraphPad Software Inc.). Statistical tests and data information are indicated in the figure legends.

## Results

### Glucose loading further attenuates BCAA catabolism in individuals with type 2 diabetes

Metabolomic profiling reveals that a glucose ingestion in fasted individuals elicits an insulin-dependent metabolic response, that is blunted with pre-diabetes [22]. To assess whether this impaired response also affects the BCAA profile, we measured levels of leucine, isoleucine, valine, and derived metabolites in plasma and *vastus lateralis* biopsies from individuals with either NGT or type 2 diabetes before and after an OGTT. Under fasted conditions, the three BCAAs were ∼10% and ∼13% higher in plasma and skeletal muscle, respectively, from individuals with type 2 diabetes as compared to NGT (Fig. 2a-c), and non-significant changes were found in the corresponding branched-chain α-keto acids (BCKAs) (Fig. 2d-f). While BCAA-derived acyl-carnitines were not different between groups (Fig. 2g-k), fasting 3-hydroxisobutyrate (3-HIB) was higher in plasma (+37%) and skeletal muscle (+45%) from individuals with type 2 diabetes as compared to NGT (Fig. 2l). This valine-derived metabolite (Fig. 2m) promotes insulin resistance in skeletal muscle cells by increasing fatty acid uptake [23].

**Figure 2.**
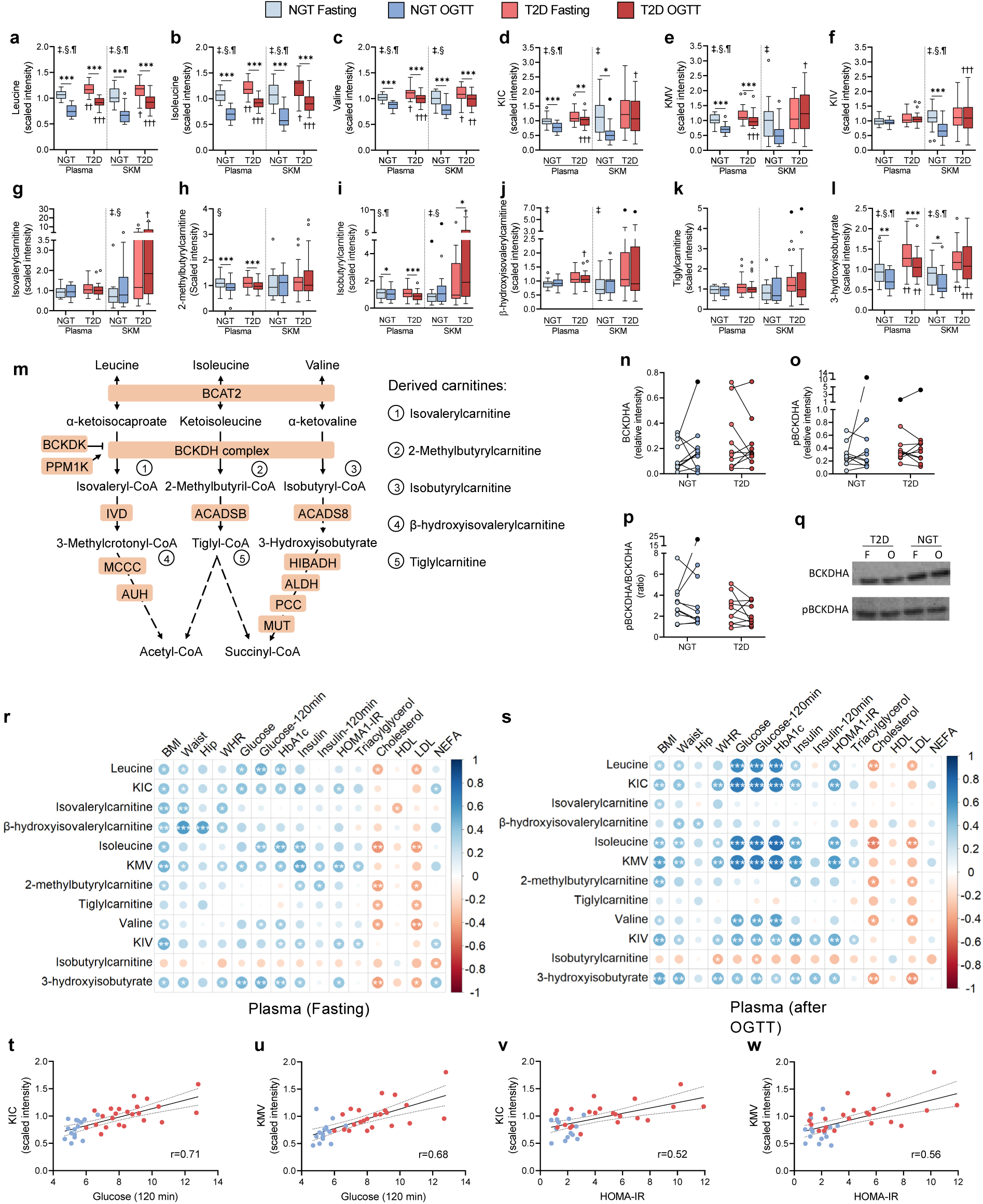
Glucose loading further attenuates BCAA catabolism in individuals with type 2 diabetes. Boxplots show scaled intensity values of BCAAs **(a-c)**, BCKAs **(d-f)**, BCAA-derived carnitines **(g-k)** and 3-hydroxisobutyrate **(l)** in plasma and skeletal muscle from individuals with NGT (n=15) or type 2 diabetes (n=26), before and after a 120-min OGTT. **m)** Catabolic pathway of BCAA and genes codifying for involved enzymes (in blue boxes). Dashed arrows indicate intermediate metabolites are not shown. **n-p)** Western blot of BCKDHA and pBCKDHA (n = 10 per group). Expression values are expressed as relative to a control sample. Black dots represent outlier values. **q)** Representatives immunoblot of the analyzed proteins in fasting conditions (F) and after an OGTT (O). **r-s)** Spearman correlation coefficients between plasma BCAA metabolites and clinical parameters. Color and size are proportional to correlation strength. **t-w)** Examples of spearman correlation coefficients summarized in panels r and s. Statistical analysis (A-P) was performed using two-way mixed-design ANOVA followed by Sidak’s *post-hoc* test. \**P*<0.05, \*\**P*<0.01, \*\*\**P*<0.001; †*P*<0.05, ††*P*<0.01, †††*P*<0.001 vs NGT. ‡, condition effect; §, glucose loading effect; ¶, interaction effect. BCAA, branched-chain amino acid; BCKA, branched-chain α-keto acid; KIC, α-ketoisocaproate; KIV, keto-isovaline; KMV, keto-methylvalerate NGT, normal glucose tolerance; SKM, skeletal muscle; T2D, type 2 diabetes. See also ESM Fig. 1.

Glucose ingestion decreased circulating and intramuscular levels of BCAA and BCKA (Fig. 2a-f, ESM Fig. 1a-b), albeit to a lesser extent in individuals with type 2 diabetes. The intramuscular content of BCKA was decreased 37–56% in individuals with NGT, whereas levels remained unaltered with type 2 diabetes. These differences were not driven by the higher BMI and age in the type 2 diabetes group (ESM Table 3). BCKA are irreversibly decarboxylated by the branched-chain α-ketoacid dehydrogenase complex (BCKDH), the rate-limiting enzyme in the catabolism of BCAA. We determined total abundance and phosphorylated (inactive) BCKDH content in skeletal muscle (Fig. 2n-q) and found no differences between groups. Glucose loading decreased 3-HIB levels in skeletal muscle of individuals with NGT, but not type 2 diabetes (Fig. 2l). Collectively, these results suggest that a glucose challenge unmasks defects at several steps of BCAA catabolism in type 2 diabetes. Indeed, circulating levels of leucine, isoleucine and derived BCKAs exhibited a positive correlation (r = 0.64 – 0.77) with blood glucose, HbA_1c_ levels and HOMA-IR after an OGTT (Fig. 2r-w, ESM Fig. 1c-d).

### Expression of genes involved in BCAA catabolism is decreased in skeletal muscle from individuals with type 2 diabetes

Expression of genes encoding for enzymes involved in the first steps of BCAA metabolism, namely Branched Chain Amino Acid Transaminase 2 (*BCAT2*) and three subunits of the BCKDH complex were decreased in skeletal muscle of individuals with type 2 diabetes (Fig. 3a). Furthermore, 69% of the analyzed genes participating in metabolic steps downstream BCKDH showed a similar profile (Fig. 3b), indicating that BCAA gene expression is widely downregulated in skeletal muscle of individuals with type 2 diabetes. These differences were also evident after an OGTT, as mRNA levels remained relatively stable. PGC-1α is an upstream regulator of BCAA metabolism [15]. Expression of *PPARGC1A*, which encodes for PGC-1α, was decreased in skeletal muscle of individuals with type 2 diabetes, irrespective of the feeding schedule (Fig. 3c). In addition, *PPARGC1A* was positively correlated with BCAA gene expression (ESM Fig. 2a) and BCAA metabolites in individuals with NGT but not type 2 diabetes (ESM Fig. 2b-c), while the expression of several of BCAA genes was inversely associated with blood glucose and HbA_1c_ (Fig. 3d). Expression of BCAA genes did not consistently correlate with BCAA-related metabolites (Fig. 3e) but exhibited opposite patterns in individuals with NGT and type 2 diabetes (ESM Fig. 2b-c).

**Figure 3.**
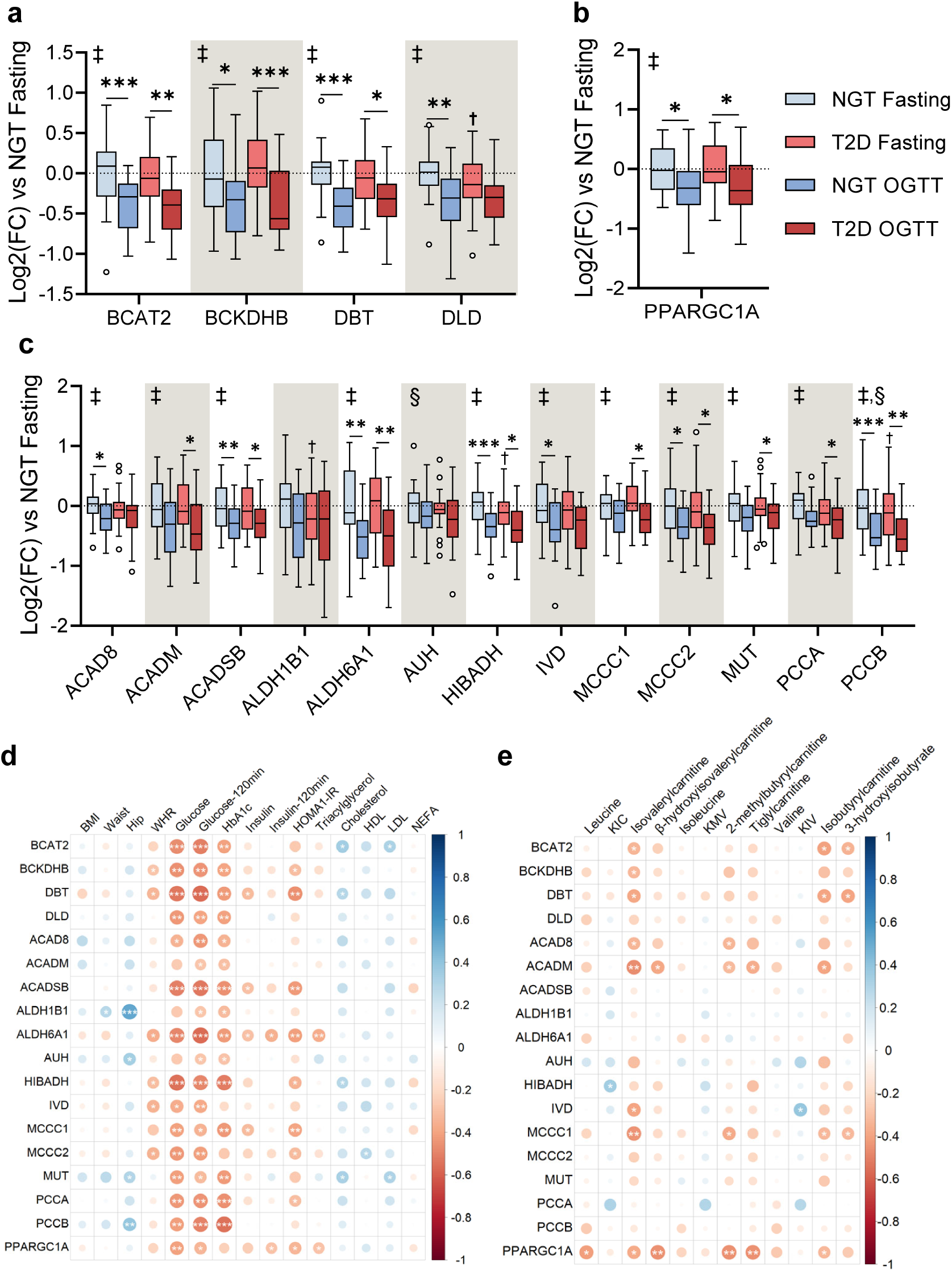
Expression of genes involved in BCAA catabolism in individuals with type 2 diabetes and correlations with blood glucose levels. **a-b)** BCAA gene expression in skeletal muscle biopsies of individuals with NGT (n=25) or type 2 diabetes (n=25) before and after a 120-min OGTT. **c)** Expression of the transcriptional coactivator *PGC-1α*. Gene expression is shown as log2(fold-change) relative to the NGT fasting group. Statistical analysis was performed using two-way mixed-design ANOVA followed by Sidak’s *post-hoc* test. **d-e)** Spearman correlation coefficients between BCAA genes and clinical parameters (d) or skeletal muscle BCAA metabolites (e) in fasted skeletal muscle. Color and size are proportional to correlation strength; \**P*<0.05, \*\**P*<0.01, \*\*\**P*<0.001; †*P*<0.05 vs fasting. ‡, condition effect; §, glucose loading effect. NGT, normal glucose tolerance; T2D, type 2 diabetes; F, fasting. See also ESM Fig. 2.

### PGC-1α mediates BCAA gene expression in primary human skeletal muscle cells

Primary HSMCs transfected with *PPARGC1A* siRNA had reduced expression of *PPARGC1A* (Fig. 4a), which moderately decreased the expression of genes of the family of acyl-CoA dehydrogenases (*ACAD8*, *ACADSB*, and *IVD*), and *HIBADH* (encoding the 3-hydroxyisobutyrate dehydrogenase enzyme), as compared to cells treated with a scrambled siRNA (Fig. 4b). Adenovirus-mediated *PPARGC1A* overexpression (Ad-PGC1A) in HSMCs (Fig. 4c) increased the expression of 61% of the genes relative to adenovirus-GFP cells (Fig. 4d). We found increased PPM1K (Fig. 4f) and reduced BCKDH protein content in Ad-PGC1A cells, which was associated with a higher pBCKDHA/BCKDHA ratio (Fig. 4g-i).

**Figure 4.**
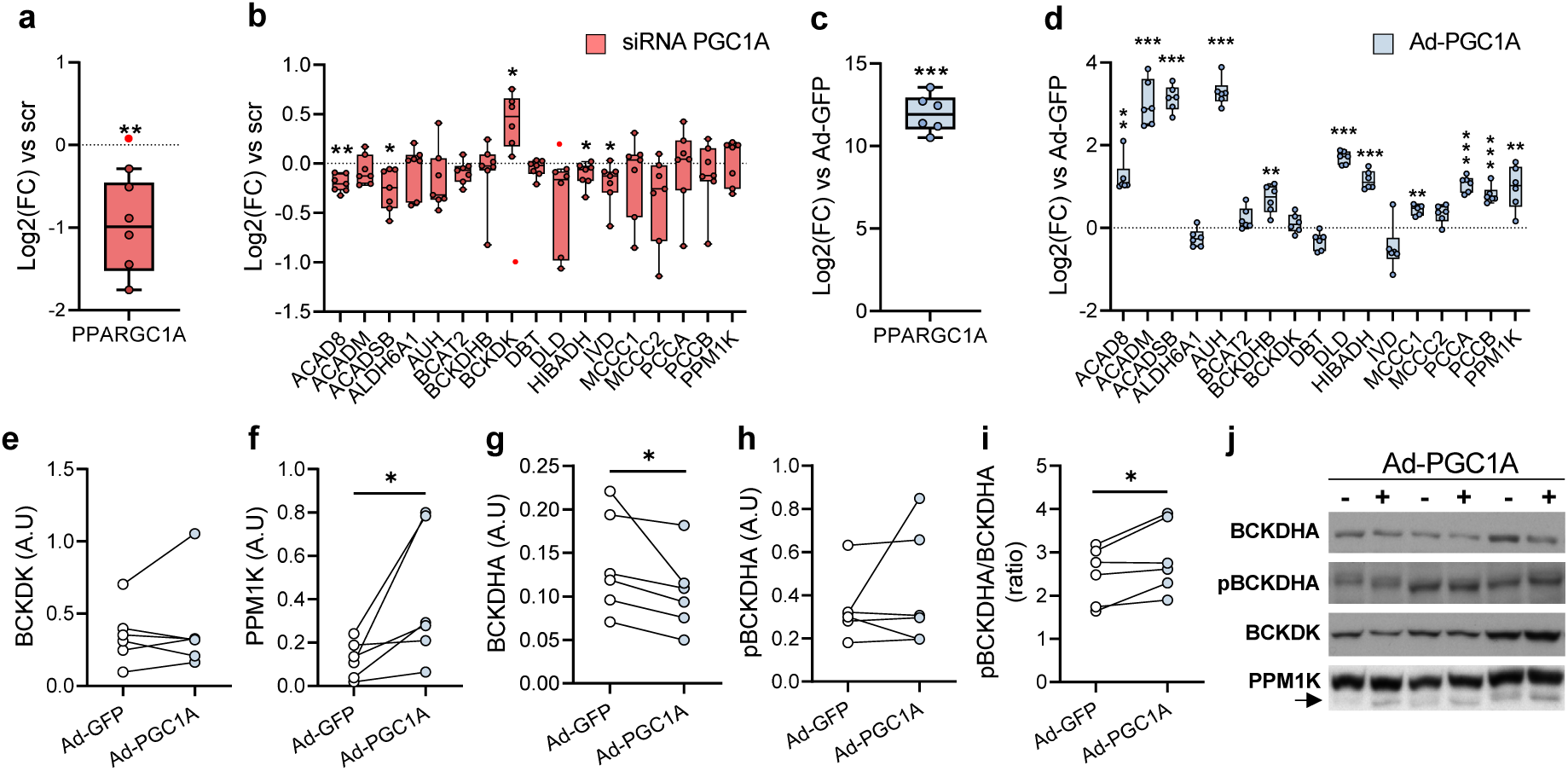
PGC-1α mediates BCAA gene expression in human skeletal muscle cells. **a)** Efficiency of *PPARGC1A* siRNA-mediated silencing. **b)** Effects of the silencing of *PPARGC1A* on the expression of BCAA catabolic genes, BCKDK and PPM1K. **c)** Expression of *PPARGC1A* in Ad-PGC1A cells. **d)** Effects of Ad-PGC1A on the expression of BCAA catabolic genes, BCKDK and PPM1K. Gene expression is shown as log2(fold-change) relative to the corresponding scramble-treated cells (dotted line). **e-i)** Protein levels of BCKDK **(e)**, PPM1K **(f),** BCKDHA **(g)**, phosphorylated BCKDHA **(h)** and ratio between pBCKDHA and BCKDHA **(i)**. **j)** Representatives immunoblot. Arrow indicates the band corresponding to PPM1K protein. Statistical analysis was performed using paired *t*-test (n=6). \**P*<0.05, \*\**P*<0.01, \*\*\**P*<0.001 vs scr (a-b) or Ad-GFP (c-i). siRNA, small interfering RNA; scr, scrambled siRNA; Ad-GFP, adenoviral overexpression of GFP; Ad-PGC1A, adenoviral overexpression of *PPARGC1A*. Red dots, significant outliers not considered in statistical calculations.

### Mice with skeletal muscle-specific modified expression of *PPARGC1A* exhibit altered levels of BCAA gene transcripts and related metabolites

Muscle-specific PGC-1α knockout mice (PGC-1α mKO) had normal body weight (ESM Fig. 3a) and slightly impaired glucose tolerance (ESM Fig. 3b-c) as compared to wild-type (WT) littermates. Skeletal muscle from PGC-1α mKO mice (Fig. 5a) had decreased expression of the majority (67%) of the analyzed BCAA genes relative to controls (Fig. 5b-d). Accordingly, a contrasting profile of BCAA gene expression was observed in skeletal muscle-specific PGC-1α transgenic mice (mTg) versus the PGC-1α mKO mice, with an upregulation of the BCAA genes relative to controls (Fig. 5e-h). These gene expression changes were not associated with alterations in either body weight or glucose tolerance (ESM Fig. 3d-f).

**Figure 5.**
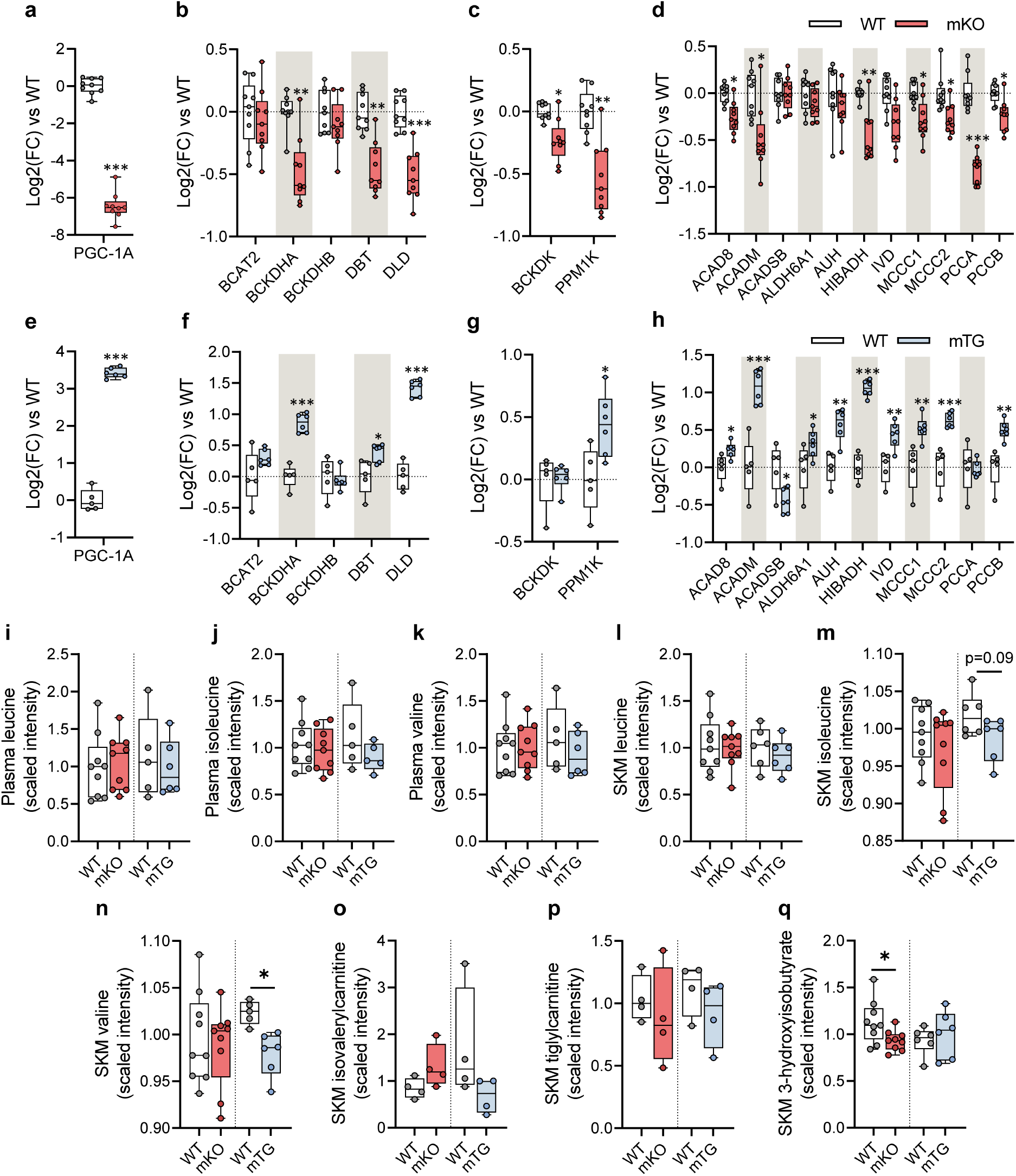
Mice with skeletal muscle-specific modified expression of PPARGC1A exhibit altered levels of BCAA gene transcripts and related metabolites. **a)** Quadriceps *PPARGC1A* expression in mKO-PGC1A mice (n=9). **b-d)** Expression of genes encoding BCAT2 and BCKDHA subunits **(b)**, BCKDK and PPM1K **(c)** and BCAA catabolic enzymes **(d)** in mKO mice. **e)** Quadriceps *PPARGC1A* expression in mTG-PGC1A mice (n=6). **f-h)** Expression of genes encoding BCAT2 and BCKDHA subunits **(f)**, BCKDK and PPM1K **(g)** and BCAA catabolic enzymes **(h)** in mTG mice. **i-k)** Scaled intensity values of leucine, isoleucine and valine in plasma from mKO, mTG and correspondent WT littermates. **l-q)** Scaled intensity values of BCAA and downstream intermediate metabolites in mKO, mTG and correspondent WT littermates. Gene expression is shown as log2(fold-change) relative to WT mice. The dotted line represents the mean of the WT group. Statistical analysis was performed using unpaired *t*-test. \**P*<0.05, \*\**P*<0.01, \*\*\**P*<0.001 vs WT. WT, wild type; mKO-PGC-1a, muscle-specific PGC-1α knockout mice; mTG-PGC-1a, muscle-specific PGC-1α transgenic mice. See also ESM Fig. S3. SKM, Skeletal Muscle.

To ascertain whether PGC-1α-associated alterations in BCAA gene expression have functional implications in BCAA metabolism, we performed LC/MS metabolomics to profile BCAA-related metabolites in plasma and *quadriceps* muscle from mKO and mTG mice. Circulating BCAA levels in mKO and mTG mice were unaltered relative to respective controls (Fig. 5i-k), whereas the intramuscular content of valine was reduced in mTG mice (Fig. 5l-n). Similar non-significant changes were observed for isoleucine (p=0.09) and isovalerylcarnitine (p=0.12) (Fig. 5m and o), while mKO mice exhibited decreased levels of 3-HIB (Fig. 5q). Consistent with our results in HSMCs, C2C12 Ad-PGC1A upregulated the expression of BCAA-related genes (ESM Fig. 3g), which was associated with increased leucine oxidation (ESM Fig. 3h).

### PGC-1α regulates BCAA gene transcription through ERRα

The orphan nuclear receptor ERRα, encoded by *ESRRA*, is a canonical functional partner of PGC-1α (Fig. 6a) that regulates metabolic processes in mitochondria [24]. Thus, we investigated whether ERRα is necessary for PGC-1α-enhanced BCAA gene expression. siRNA-mediated silencing of ERRα (Fig. 6b) decreased the expression of 65% of the analyzed genes (Fig. 6c). We determined whether this BCAA gene downregulation affects leucine oxidation in skeletal muscle. HSMCs were treated with either DMSO or BT2, a BCKDK inhibitor that increases BCAA oxidative flux. *ESRRA* silencing decreased leucine oxidation rates under BCKDH-activated conditions, as measured by CO_2_ production after incubation with [U–^14^C]-leucine (Fig. 6d). We next tested whether ERRα is a PGC-1α interacting partner in the transcriptional regulation of the BCAA gene set using HSMCs treated with either scramble siRNA or *ESRRA* siRNA and transfected with Ad-GFP or Ad-PGC1A. We also tested this using Ad-PGC-1A cells treated with the inverse ERRα agonist XCT-790 (ESM Fig. 4). Since *PPARGC1A* and *ESRRA* mutually regulate their expression, we confirmed that *ESRRA* silencing did not abrogate *PPARGC1A* in cultured cells. Ad-PGC1A cells had high levels of *PPARGC1A* transcripts regardless the treatment with *ESRRA* siRNA (Fig. 6e and ESM Fig. 4a), whereas the expression of two known targets of PGC-1α/ERRα, *TFAM* and *VEGF*, was dampened by *ESRRA* siRNA despite the overexpression of *PPARGC1A*. Knockdown and inhibition of *ESRRA* completely ablated PGC-1α-mediated upregulation in all analyzed genes (Fig. 6f-h, ESM Fig. 4b-d), indicating that ERRα is essential for the transcriptomic regulation of the BCAA gene network orchestrated by PGC-1α. Nevertheless, *ESRRA* expression was similar between fasted individuals with either NGT or type 2 diabetes (Fig. 6i).

**Figure 6.**
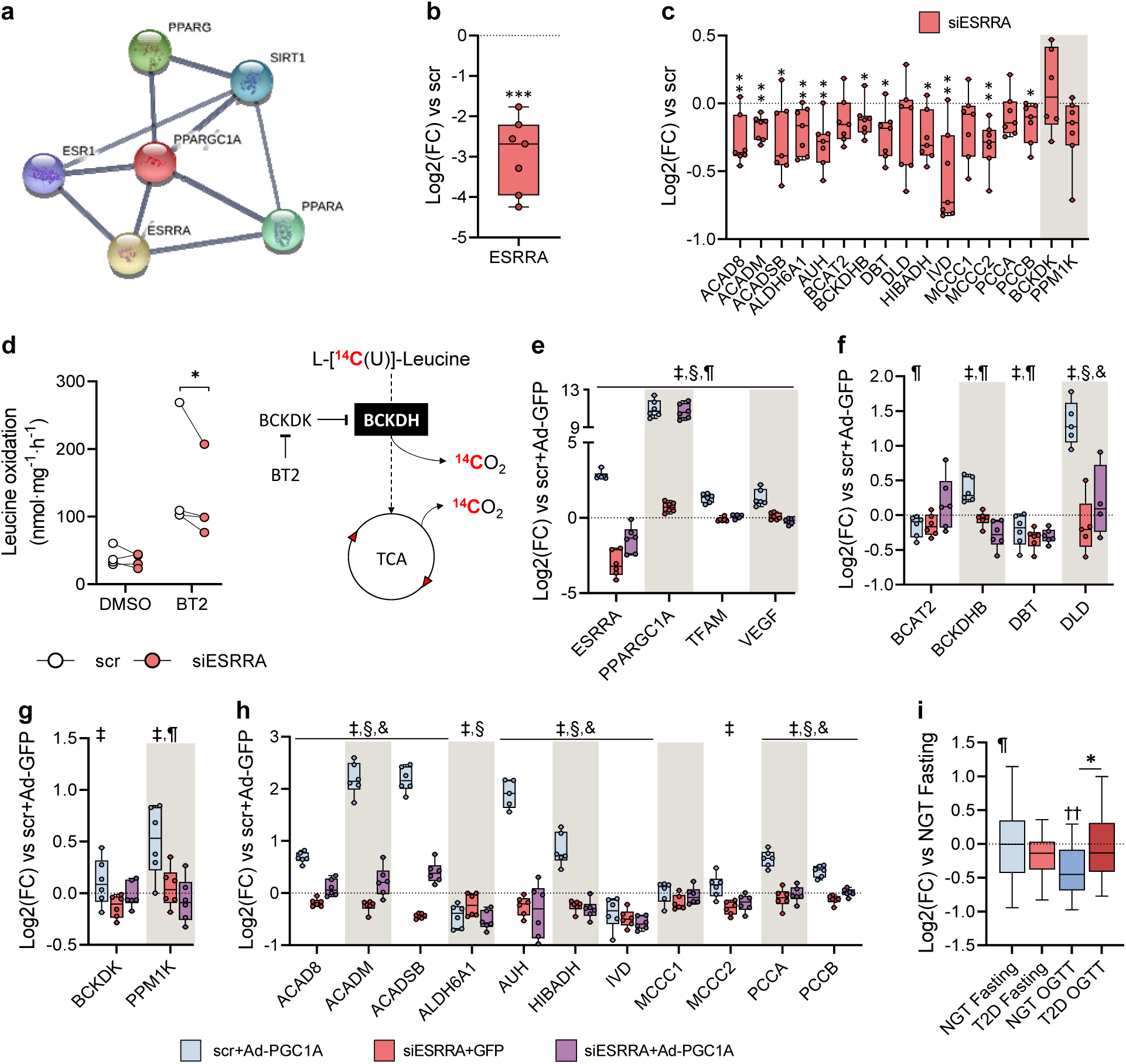
PGC-1α regulates BCAA gene transcription through ERRα. **a)** STRING Interacting Network showing top 5 interacting proteins for PGC-1α. **b)** Efficiency of ERRα silencing in primary HSMCs. **c)** Expression of BCAA genes. **d)** Leucine oxidation rates in HSMCs. Schematic biochemical representation of the leucine oxidation assay is shown. **e)** Expression of *ESRRA*, *PPARGC1A* and target genes in Ad-PGC1A cells transfected with *ESRRA* siRNA. **f-h)** BCAA gene expression in Ad-PGC-1A cells with or without *ESRRA* siRNA. **i)** Expression of *ESRRA* in skeletal muscle biopsies from individuals with NGT or type 2 diabetes. Gene expression is shown as log2(fold-change) relative to the corresponding scramble+GFP treated cells (dotted line) scr (b-c), scr+Ad-GFP (E-H) or NGT Fasting (i). Statistical analysis was performed using paired *t*-test (a-b, n=7), two-way repeated measures ANOVA followed by Tukey’s *post-hoc* test (d, n=4; e-h, n=6), or two-way mixed-design ANOVA followed by Sidak’s post-hoc test (i, n=25). \**P*<0.05, \*\**P*<0.01, \*\*\**P*<0.001 vs scr (a-b) or scr-Ad-GFP (c-f). ‡, siESRRA effect; §, Ad-PGC1A effect; ¶, interaction effect. siRNA, small interfering RNA; Ad-GFP, adenoviral overexpression of GFP; Ad-PGC1A, adenoviral overexpression of *PPARGC1A*; BT2, 3,6-dichlorobrenzo(b)thiophene-2carboxylic acid; scr, scrambled siRNA; TCA, tricarboxylic acid cycle. See also ESM Fig. 4.

## Discussion

A link between increased plasma BCAAs and insulin resistance was established as early as 1969 [3], but this association has remained relatively unexplored until the last decade. Here we elucidate the mechanisms underpinning BCAA metabolism in type 2 diabetes. While we corroborate the association between BCAA metabolites and type 2 diabetes [9, 25–27] and the role of PGC-1α in BCAA catabolism [15, 23, 28], we provide new evidence that an OGTT unmasks impairments in BCAA catabolism in individuals with type 2 diabetes. Moreover, we reveal that PGC-1α-mediated regulation of genes important for BCAA catabolism is dependent on ERRα, a canonical PGC-1α-interacting protein.

Circulating plasma BCAA levels are altered in individuals with type 2 diabetes, however metabolomic profiles of peripheral tissues involved in glucose homeostasis are scarce and the transcriptomic regulation of genes involved in BCAA catabolism is unknown. Studies focusing on skeletal muscle BCAA content are limited to an analysis of adults with insulin resistance [14], rather than type 2 diabetes. We performed untargeted metabolomic analysis on both plasma and *vastus lateralis* biopsies obtained before and after an OGTT from men with either NGT or type 2 diabetes. Corroborating an earlier study of plasma samples from the Framingham Heart Study Offspring cohort [29], circulating BCAAs were decreased after an OGTT and this reduction was attenuated in type 2 diabetes. Concomitantly, we found that this blunted response also affected leucine- and isoleucine-derived BCKAs. In skeletal muscle, insulin inhibits proteolysis [30] and increases the preference for BCAA oxidation [13], which may explain the blunted excursion of BCAA and BCKA metabolites in insulin-resistant individuals with type 2 diabetes as compared to NGT. Consistent with this hypothesis, BCAAs, BCKAs and 3-HIB levels after the glucose challenge were significantly less decreased in the skeletal muscle from individuals with type 2 diabetes. Although we did not detect changes in BCKDHA phosphorylation state, BCAA catabolic flux cannot be predicted by BCKDH phosphorylation status [13], thus, we cannot exclude the possibility that BCKDH activity *per se* is impaired. Moreover, insulin modulates BCAA transamination in type 2 diabetes [31] and BCAT2 activity [32], suggesting that insulin resistance could lead to an accumulation of BCKAs due to defects in both BCAA transamination and oxidation.

Generally, plasma metabolites mirrored skeletal muscle BCAA profile. However, the intramuscular accumulation of isovalerylcarnitine, isobutyrylcarnitine, and 3-HIB was not reflected in plasma, suggesting defects in processes controlling the export of metabolites from myocytes in type 2 diabetes. Accordingly, the accumulation of 3-HIB after an OGTT is notable, since this valine-derived metabolite promotes insulin resistance through increased fatty acid uptake in skeletal muscle [23]. This could lead to a vicious cycle in which secretion of 3-HIB from skeletal muscle may decrease insulin sensitivity and further impair insulin signaling. Although elevated levels of circulating C3 and C5 acylcarnitines have been also detected in individuals with obesity [7], whether accumulation of short-chain acylcarnitines mediates in insulin resistance remains to be elucidated [33]. These results suggest the degradation of BCAA-derived metabolites is incomplete. Additionally, we found a strong correlation of post-OGTT BCAAs and derived BCKAs circulating levels with blood glucose and HbA_1c_. Since plasma fasting BCAA levels exhibited much weaker relationship with clinical biomarkers of type 2 diabetes, this finding underscores a tight connection between impaired glucose homeostasis and whole-body BCAA catabolism.

The accumulation of BCAA metabolites in individuals with type 2 diabetes was accompanied by reduced expression of genes involved in BCAA catabolism, suggesting that alterations in the transcriptional regulation of these genes could attenuate BCAA catabolism in skeletal muscle. Contrasting a previous report [34], we did not find a correlation between BCAA genes and related metabolites, suggesting that posttranscriptional modifications play a key role in the impairment of BCAA catabolism. Conversely, the BCAA gene set was inversely correlated with blood glucose levels, suggesting a connection between glucose homeostasis and alterations in the transcriptional regulation of genes involved in BCAA catabolism.

PGC-1α mediates the expression of genes involved in BCAA catabolism [15, 23, 28]. Concordantly, expression of *PPARGC1A* was positively correlated with BCAA genes. PGC-1α also coordinates metabolic and transcriptomic programs linked to cellular energy homeostasis [35–37], and reduced PGC-1α mRNA and protein levels are linked to insulin resistance in type 2 diabetes [38, 39]. In the present study, primary HSMCs and mouse skeletal muscle with reduced *PPARGC1A* expression exhibited a mild reduction in expression of several BCAA genes, whereas PGC-1α overexpression was associated with a consistent upregulation. Thus, while PGC-1α was not essential for basal BCAA gene transcription, a role in the adaptive response to increased BCAA catabolic demands, such as during exercise or nutritional changes cannot be excluded [40]. We next hypothesized that alterations in the BCAA gene network would impact BCAA metabolism. We found that mice overexpressing PGC-1α in skeletal muscle exhibited an upregulation of BCAA genes and reduced intramuscular accumulation of valine, suggestive of increased BCAA catabolic flux. Similarly, adenovirus-mediated overexpression of PGC-1α in C2C12 myotubes increased leucine oxidation rates. However, intramuscular levels of BCAA metabolites were unaltered in PGC-1α mKO mice. Thus, other organs such as adipose tissue may compensate for a putative impairment in BCAA catabolism [41]. Furthermore, a metabolic challenge may be necessary to reveal functional alterations in BCAA metabolism in PGC-1α mKO and mTG mice.

PGC-1α is a transcriptional coactivator and does not directly bind DNA in a sequence-specific manner. The nuclear orphan receptor ERRα is one of the main transcriptional partners of PGC-1α [24], and through this physically interaction, elicits the transcription of genes involved in mitochondrial biogenesis and oxidative phosphorylation [42], lipid metabolism [43] and ketone body homeostasis [44]. We found that *ESRRA* silencing, as well as inhibition of ERRα with an inverse agonist, downregulated BCAA genes and abrogated the PGC-1α-induced responses, indicating ERRα is necessary for the PGC-1α-mediated transcription of BCAA genes. Moreover, *ESRRA* silencing was associated with a modest decrease in BCKDH-stimulated leucine oxidation, suggesting a functional impact of the transcriptional downregulation of BCAA genes under energy demanding conditions. Thus, we next hypothesized that alterations in ERRα may explain the defects in BCAA catabolism in the individuals with type 2 diabetes. However, expression of *ESRRA* was unaltered in skeletal muscle of individuals with NGT or type 2 diabetes, suggesting that reductions in PGC-1α is sufficient to impair BCAA expression or alternatively, the PGC-1α/ERRα interaction is compromised in type 2 diabetes.

Some limitations of our study warrant discussion. Our analysis was limited to male participants, and therefore we cannot exclude sex-specific differences in the analyzed outcomes. Most of the individuals with type 2 diabetes included in this study were treated with metformin. Considering mitochondria are the major target of metformin, this may affect BCAA metabolism. Due to limitations in the LC/MS methodology, quantitation of BCAA metabolites with CoA moieties was not possible. To overcome this, we used specific derived carnitines from these metabolites as a proxy. Alterations in PGC-1α expression may influence fiber type distribution [19, 45], which could impact the results. Nevertheless, similar results were obtained in transgenic mice, *in vitro* primary HSMCs and C2C12 myotubes, suggesting that alterations in BCAA metabolism are unrelated to fiber type.

In conclusion, altered expression of *PPARGC1A* is associated with disturbances in BCAA metabolism in skeletal muscle of individuals with type 2 diabetes. Experimental approaches to reduce *PPARGC1A* levels partially recapitulates the BCAA gene set profile identified in skeletal muscle from type 2 diabetes patients, without impacting circulating and intramuscular BCAA levels. Our results indicate that glucose loading exacerbates disturbances in the BCAA profile, revealing that the metabolic inflexibility that characterizes type 2 diabetes encompasses BCAA catabolism. Additionally, our data demonstrate that ERRα is essential for PGC-1α-mediated transcriptional regulation of genes involved in BCAA metabolism in primary human myotubes, thereby unraveling a new role for this orphan nuclear receptor.

## Abbreviations

3-HIB: 3-hydroxyisobutyrate
BCAA: branched-chain amino acid
BCKA: branched-chain α-keto acid
ERRα: Estrogen-related receptor alpha
HSMC: human skeletal muscle primary cell
NGT: normal glucose tolerance
PGC1α: Peroxisome proliferator-activated receptor gamma coactivator 1-alpha
WT: Wild type

## Data availability

The datasets generated during and/or analyzed during the current study are available from the corresponding author on reasonable request.

## Author Contributions

R.J.O.S. conceived the idea, planned the experiments and collected and analyzed data. D.R.R performed leucine oxidation assays, analyzed data and wrote the manuscript. A.V.C. contributed to discussion and interpretation of metabolomic and protein data. E.C. collected mouse metabolite data and contributed to discussion. R.F. assisted with animal care, OGTT and skeletal muscle sampling. S.K. contributed to the conception of the idea. J.H. contributed to the conception of the idea. H.K.K. assisted with recruitment and collection of human metabolite data. C.H. provided mouse biological samples and contributed to the discussion of animal data. T.M. supervised acquisition and analysis of mouse metabolite data. A.K. supervised the study and contributed to the discussion of all results. E.N. assisted with recruitment and obtained human skeletal muscle biopsies and blood samples. J.R.Z. supervised the study, edited the manuscript and acquired funding. All authors have reviewed the article and gave their final approval. J.R.Z is the guarantor of this work and, as such, had full access to all the data in the study and takes responsibility for the integrity of the data and the accuracy of the data analysis.

## Electronic Supplementary Materials (ESM)

### ESM Methods

#### Human plasma and skeletal muscle metabolomics

Metabolomic analysis was performed by Metabolon, Inc. (Durham, NC, USA) as described (20). Equal volume of plasma and equal weights of *vastus lateralis* biopsy were used for all samples. The protein fraction was removed using a methanol extraction method while allowing maximum recovery of small molecules. The resulting extract was divided into five fractions: one for analysis by ultra-high-performance liquid chromatography-tandem mass spectroscopy (UPLC-MS/MS) with positive ion mode electrospray ionization, one for UPLC-MS/MS with negative ion mode electrospray ionization, one for liquid chromatography (LC) polar platform, one for gas chromatography–MS (GC–MS), and one sample was reserved as a backup.

The LC/MS portion of the UPLC-MS/MS platform was based on a Waters ACQUITY UPLC and a Thermo Scientific Q-Exactive high resolution/accurate mass spectrometer interfaced with a heated electrospray ionization source and Orbitrap mass analyzer operated at 35,000 mass resolution. Separated dedicated columns (Waters UPLC BEH C18-2.1×100 mm, 1.7 µm) were used to perform analysis under acidic positive ion optimized conditions and basic negative ion optimized conditions. The third aliquot was analyzed via negative ionization following elution from a HILIC column (Waters UPLC BEH Amide 2.1×150 mm, 1.7 µm) using a gradient consisting of water and acetonitrile with 10 mM ammonium formate.

Samples destined for GC-MS analysis were dried under vacuum for at least 18 h prior to being derivatized under dried nitrogen using bistrimethyl-silyltrifluoroacetamide. Derivatized samples were separated on a 5% diphenyl / 95% dimethyl polysiloxane fused silica column (20 m x 0.18 mm ID; 0.18 µm film thickness) with helium as carrier gas and a temperature ramp from 60°C to 340°C within a 17.5-min period. Samples were analyzed on a Thermo-Finnigan Trace DSQ fast-scanning single-quadrupole mass spectrometer using electron impact ionization and operated at unit mass resolving power. The scan range was from 50–750 m/z.

Compounds were identified by comparison to library entries of purified standards or recurrent unknown entities. Metabolon maintains a library based on authenticated standards that contains the retention time/index (RI), mass to charge ratio (m/z), and chromatographic data on all molecules present in the library. Biochemical identifications were based on three criteria: retention index within a narrow RI window of the proposed identification, accurate mass match to the library and the MS/MS forward and reverse scores between the experimental data and authentic standards. The MS/MS scores were based on a comparison of the ions present in the experimental spectrum to the ions present in the library spectrum.

#### Mouse plasma and skeletal muscle metabolomics

Mouse sample preparation and analysis was performed at the Swedish Metabolomics Centre (Umeå University). Prior to gas chromatography time-of-flight mass spectrometry (GC-TOF/MS) analysis, a two-step derivatization procedure was carried out to increase metabolites volatility and reduce number of tautomeric forms. Derivatized samples were analyzed on an Agilent 6890 gas chromatograph equipped with a 10 m × 0.18 mm i.d. fused silica capillary column with a chemically bonded 0.18-µm DB 5-MS stationary phase (J&W Scientific, Folsom, CA, USA). The injector temperature was 270°C. The column temperature was held at 70°C for 2 min, increased by 40°C min^-1^ to 320°C, and held there for 1 min. The column effluent was introduced into the ion source of a Pegasus III time-of-flight mass spectrometer, GC-TOF/MS (Leco, St. Joseph, MI, USA). The transfer line and ion source temperatures were 250°C and 200°C, respectively. Ions were generated by a 70-eV electron beam at an ionization current of 2.0mA. Spectra were recorded in the mass range 50–800 m/z at 30 spectra s^-1^.

Samples destined to liquid chromatography time-of-flight mass spectrometry (LC-TOF/MS) were re-suspended in methanol and water (1:1) and injected onto a Waters Acquity UPLC HSS T3 C18 column (2.1 × 50 mm, 1.8 μm; Waters Corporation, Milford, MA, USA) in combination with a 2.1 mm x 5 mm, 1.8 µm VanGuard precolumn (Waters Corporation) held at 40°C. Metabolites chromatographic separation were carried out using a gradient solvent system consisting of water with 0.1% formic acid and acetonitrile/isopropanol (75/25, v/v) with 0.1% formic acid at a flow rate of 0.5 mL min^-1^. The detection of separated metabolites was performed using the Agilent 6550 Q-TOF mass spectrometer equipped with a jet stream electrospray ionization source, operating in both positive and negative ion modes. A reference interface was connected for accurate mass measurements; the reference ions purine (4 µM) and HP-0921 (Hexakis (1H, 1H, 3H-tetrafluoropropoxy) phosphazine) (1 µM) were infused directly into the MS at a flow rate of 0.05 mL min-1 for internal calibration. Full scan MS spectra were collected in a centroid mode over the mass range 70-1700 m/z with an acquisition rate of 4 spectra s^-1^. The Auto MS/MS was performed on QC-samples. The isolation width was set to narrow, i.e. approx. 1.3 amu, the mass range was 40-1700, and data was collected in centroid mode with an acquisition rate of 3 scans s-1. The collision energy was 10, 20 and 40 ev. Data were acquired with MassHunter Acquisition Software B.07.01.

The processing of GC-TOF/MS data and extraction of putative metabolites was conducted as described (21). Briefly, an in-house MATLAB script was used for the extraction of putative metabolites by matching the mass spectra and retention indices to in-house mass spectral library at the Swedish Metabolomics Centre and the publicly available Max Planck Institute library in Golm. The processing of LC-TOF/MS data and extraction of putative metabolites and lipids were performed by MassHunter Profinder version B.08.00 in combination with Qualitative Analysis software version B.07.00, PCDL manager version B.07.00 and Mass Profiler Professional™ 13.0 (all from Agilent Technologies Inc., Santa Clara, CA, USA). Annotation of metabolites were done by matching the retention time (MS and MS-MS spectra) against the in-house metabolite library.

#### RNA isolation and relative mRNA expression

Total RNA from human skeletal muscle biopsies, mouse skeletal muscles and cultured myotubes was extracted using TRIzol according to manufacturer’s instructions (Thermo Fisher Scientific). Concentration and extraction quality (A260/A280) of RNA samples were determined by spectrophotometry using a Nanodrop ND-1000 (Thermo Fisher Scientific) and equal amounts of RNA were used for cDNA synthesis. cDNA was synthesized using the High Capacity cDNA Reverse Transcription kit with random primers (4368814, Thermo Fisher Scientific). RT-qPCR was performed using either a ViiA 7 Real-Time PCR 384-well system or a StepOne Plus Real-Time PCR 96-well system (Thermo Fisher Scientific). Gene expression was determined using Fast SYBR™ Green Master Mix (4309155, Thermo Fisher Scientific). Sequences of primers used for the real-time PCR are reported in ESM Tables 1 and 2. Gene expression in human skeletal muscle biopsies and cultured myotubes was normalized to the geometrical mean of *GUSB*, *RPLP0* and *TBP* expression. For C2C12 cells and mouse tibialis anterior muscle, gene expression was normalized to the geometrical mean of *B2M*, *RPLP0* and *TBP* expression. Relative gene expression was calculated by the comparative ΔΔCt method and normalized to control group (in human and mouse samples, NGT Fasting and WT mice, respectively) or control samples (*in vitro* experiments, cells from each donor were normalized against their correspondent siRNA scr-treated counterparts).

#### Western blot analysis

Human skeletal muscle biopsies were homogenized in ice-cold lysis buffer [137 mM NaCl, 2.7 mM KCl, 1 mM MgCl2, 0.5 mM Na_3_VO_4_, 10% (vol/vol) glycerol, 1% (vol/vol) Triton X-100, 20 mM Tris, 10 mM NaF, 1 mM EDTA, 1 mM PMSF, and 1% (vol/vol) protease inhibitor cocktail set 1 (Merck, Darmstadt, Germany)] using a TissueLyser II (21 Hz, 45 sec, two times. Qiagen, Haan, Germany). Homogenates were rotated for 30 min at 4°C and thereafter subjected to centrifugation at 12000 g for 15 min at 4°C. Cells were lysed in ice-cold homogenization buffer. Lysates were rotated for 30 min at 4°C and subjected to centrifugation at 12000 g for 15 min at 4°C.

Protein concentration in the supernatants of skeletal muscle homogenates and cell lysates was determined by BCA protein assay kit (Pierce Biotechnology, Rockford, IL, USA). Samples were prepared with Laemmli buffer to equal final protein concentrations, separated on Criterion XT Bis-Tris Gels for SDS-PAGE (Bio-Rad, Hercules, CA; USA) and transferred to PVDF membranes (Merck). Thereafter, membranes were stained with Ponceau S to verify transfer quality and confirm equal protein loading. After blocking with 7.5% nonfat milk in Tris-buffered saline with Tween (TBST; 10 mM Tris-HCl, 100 mM NaCl, 0.02% Tween 20) for 2 h at room temperature, membranes were incubated overnight at 4°C with primary antibodies against β-actin (A5441, Sigma-Aldrich), GAPDH (sc-47724, Santa Cruz Biotechnology), BCKDHA (ab68094, Abcam), p^S293^-BCKDHA (ab200577, Abcam), BCKDK (sc-374425, Santa Cruz Biotechnology), and PPM1K (ab67935, Abcam), all at a 1:1000 dilution. Antibodies were validated in HSMCs in which the corresponding encoding gene was silenced with siRNA. Membranes were washed with TBST and incubated with horseradish peroxidase-conjugated secondary antibodies (1:10000 – 1:25000). Proteins were visualized by enhanced chemiluminescence (Amersham ECL Western Blotting Detection Reagent, Little Chalfont, UK). Bands of interest were semi-quantified by densitometry (QuantityOne, Bio-Rad).

## ESM Tables

**ESM Table 1.**
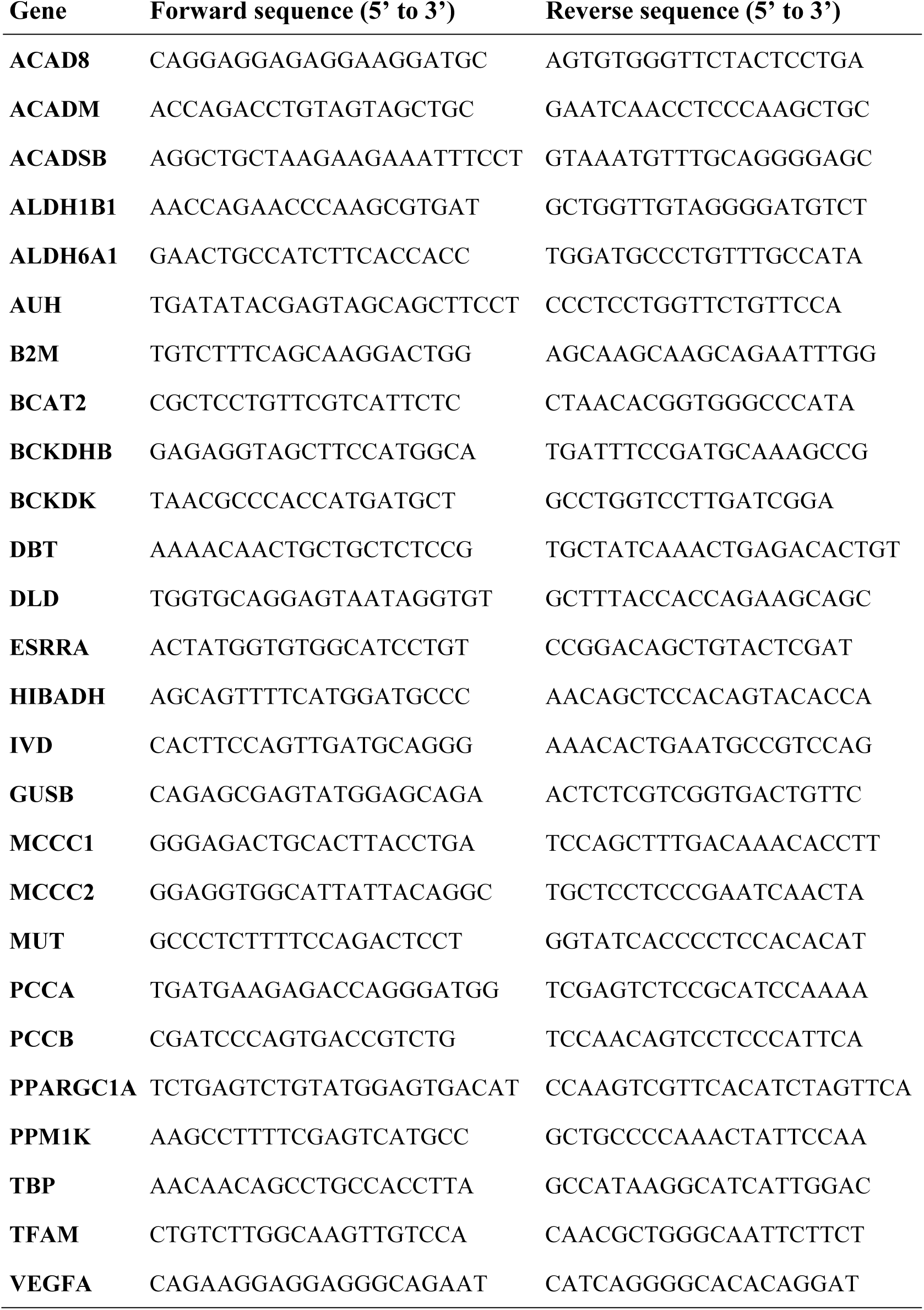
Forward and reverse sequences of primers used in gene expression experiment analysis of human skeletal muscle.

**ESM Table 2.**
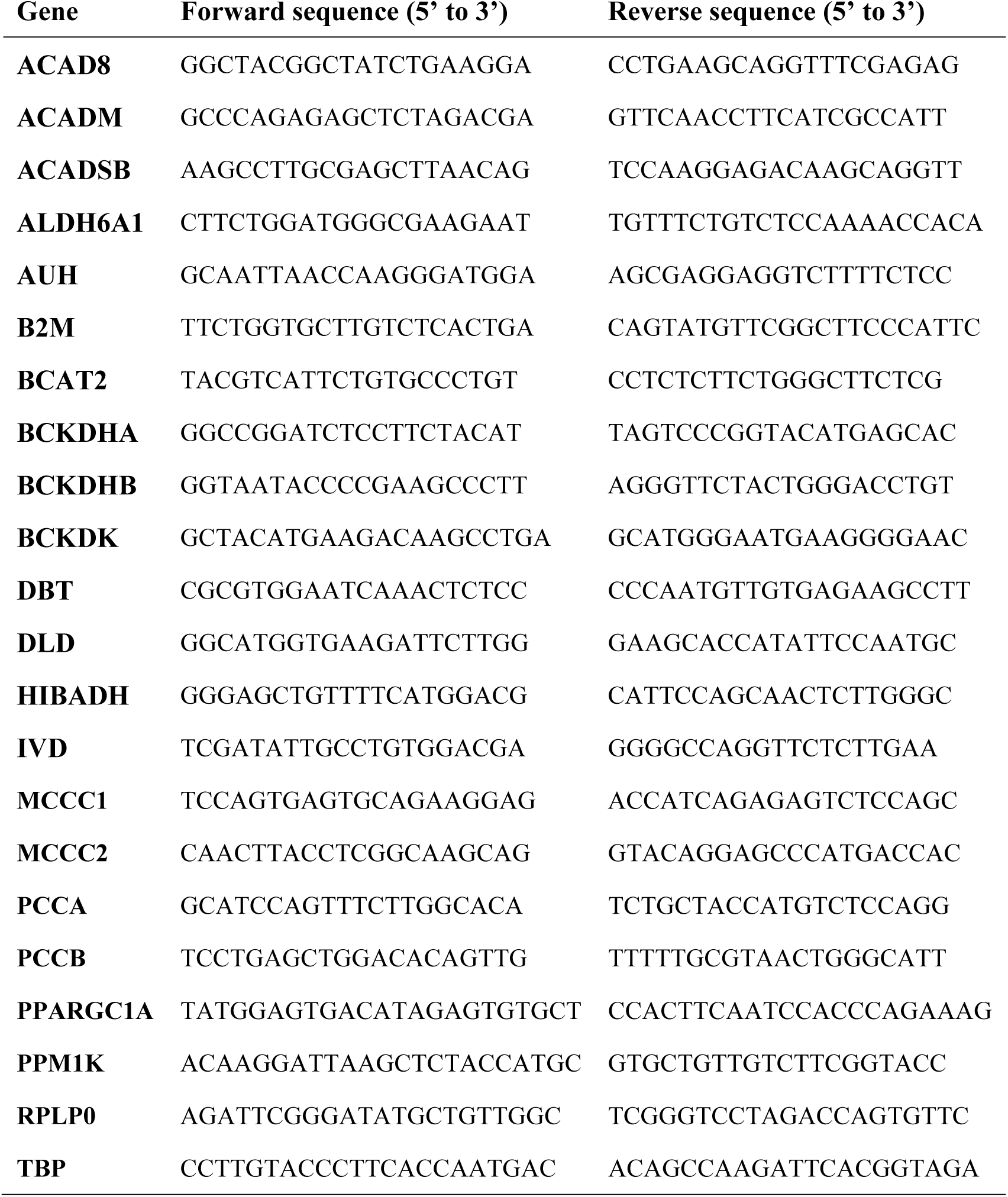
Forward and reverse sequences of primers used in gene expression analysis of mouse skeletal muscle.

**ESM Table 3.** Y-intercept p-values corresponding to the analysis of covariance comparing individuals with normal glucose tolerance and type 2 diabetes controlling for variation in BMI and age.

|  | BMI |  | Age |  |
| --- | --- | --- | --- | --- |
|  | Plasma | SKM | Plasma | SKM |
| **Leucine** | <0.0001 | 0.0008 | <0.0001 | <0.0001 |
| **Isoleucine** | <0.0001 | 0.0008 | <0.0001 | <0.0001 |
| **Valine** | 0.0013 | 0.0092 | 0.0003 | 0.0006 |
| **α-ketoisocaproate** | 0.0014 | 0.0300 | 0.0002 | 0.0024 |
| **Ketoisoleucine** | 0.0021 | 0.0175 | 0.0403 | 0.0106 |
| **α-ketovaline** | 0.1540 | 0.0184 | 0.0431 | 0.0011 |
| **Isovalerylcarnitine** | 0.4343 | 0.0166 | 0.1120 | 0.0199 |
| **2-methylbutyrylcarnitine** | 0.9525 | 0.1697 | 0.1535 | 0.1250 |
| **Isobutyrylcarnitine** | 0.5386 | 0.0244 | 0.4478 | 0.0189 |
| **β-hydroxyisovalerylcarnitine** | 0.3515 | 0.0625 | 0.3134 | 0.0370 |
| **Tiglylcarnitine** | 0.8905 | 0.1711 | 0.5280 | 0.2819 |
| **3-hydroxyisobutyrate** | 0.0132 | 0.0116 | 0.0008 | 0.0006 |

Linear regression between BCAA metabolites and BMI/Age was calculated in the two experimental groups (after OGTT). Obtained slopes were in all cases not statistically different between groups, allowing to compare Y-intercepts of the regression lines.

## ESM Figures

**ESM Figure 1 (Supplementary to Figure 2).**
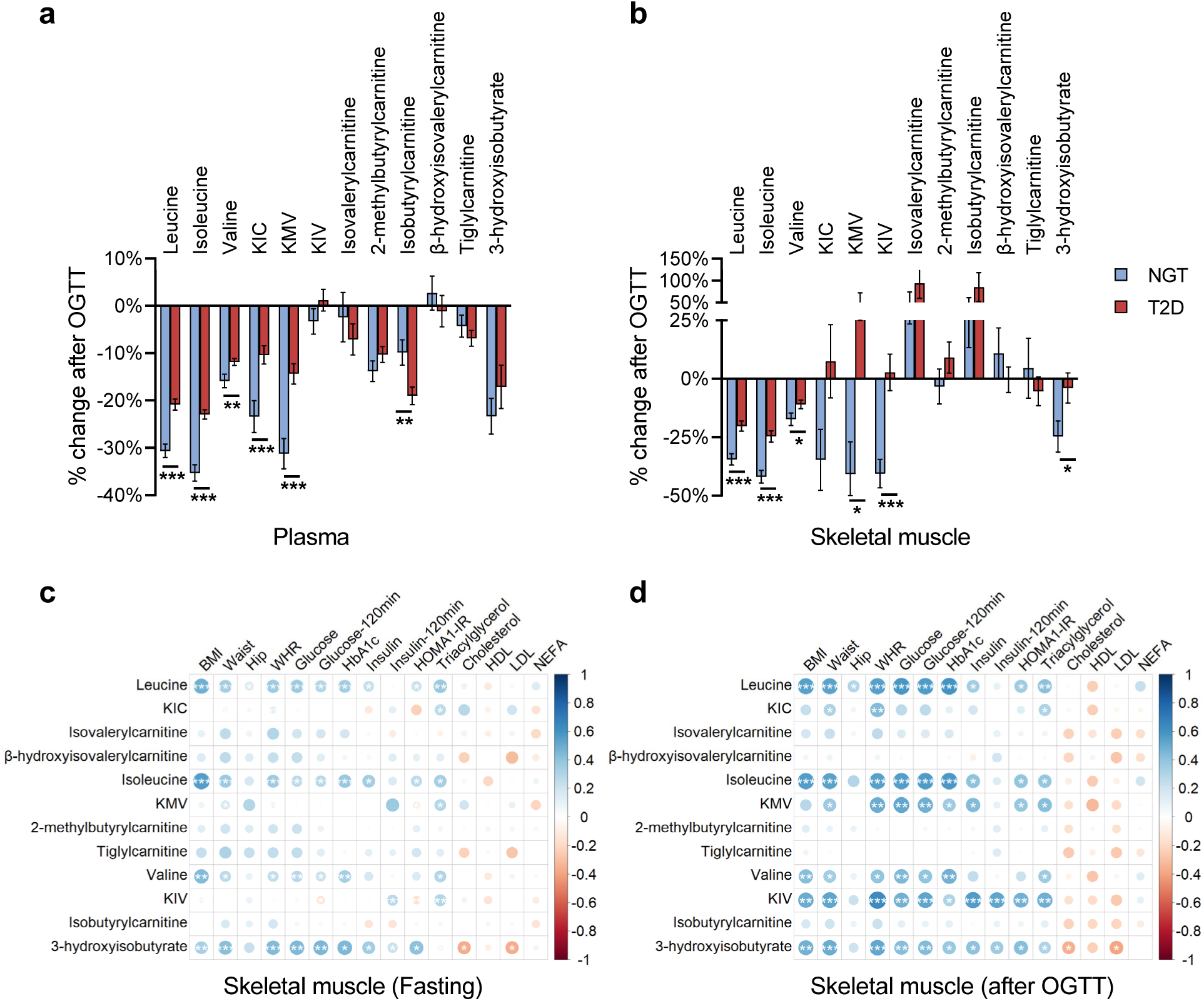
**a-b)** Metabolite excursions after OGTT in individuals with normal glucose tolerance (NGT) and type 2 diabetes (T2D). **c-d)** Spearman correlation coefficients between clinical parameters and skeletal muscle BCAA metabolites measured before (**c**) and after (**d**) and OGTT. Color and size are proportional to correlation strength; *, *P*<0.05, **, *P*<0.01 and ***, *P*<0.001. KIC, α-ketoisocaproate; KIV, keto-isovaline; KMV, keto-methylvalerate.

**ESM Figure 2 (Supplementary to Figure 2).**
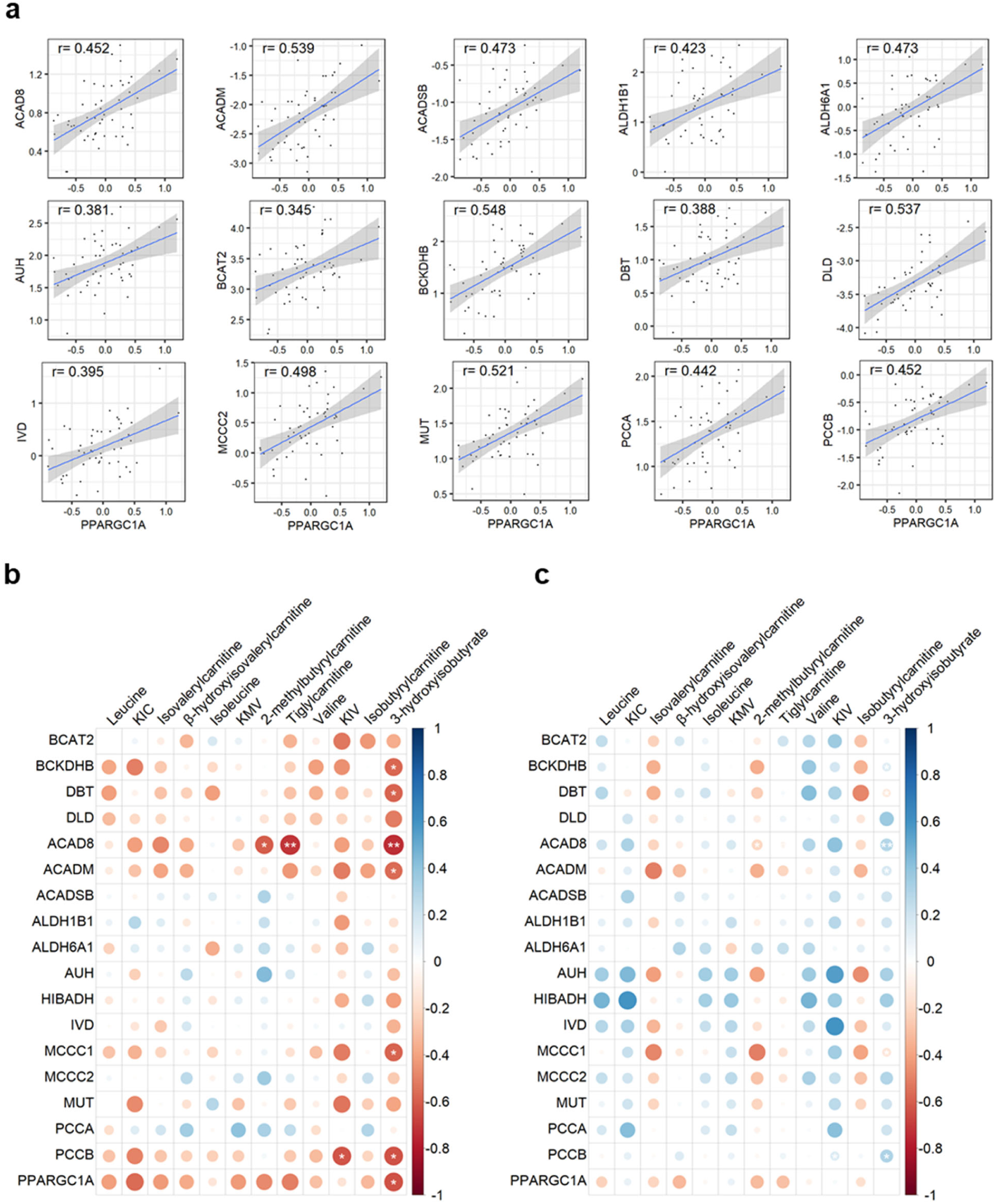
**a)** Spearman correlation between the skeletal muscle expression of *PPARGC1A* and BCAA genes. **b-c)** Spearman correlation coefficients between expression of genes involved in BCAA catabolism and skeletal muscle BCAA metabolites measured in individuals with normal glucose tolerance (**b**) and with type 2 diabetes (**C**). Color and size are proportional to correlation strength; *, *P*<0.05, **, *P*<0.01 and ***, *P*<0.001. KIC, α-ketoisocaproate; KIV, keto-isovaline; KMV, keto-methylvalerate.

**ESM Figure 3 (Supplementary to Figure 5).**
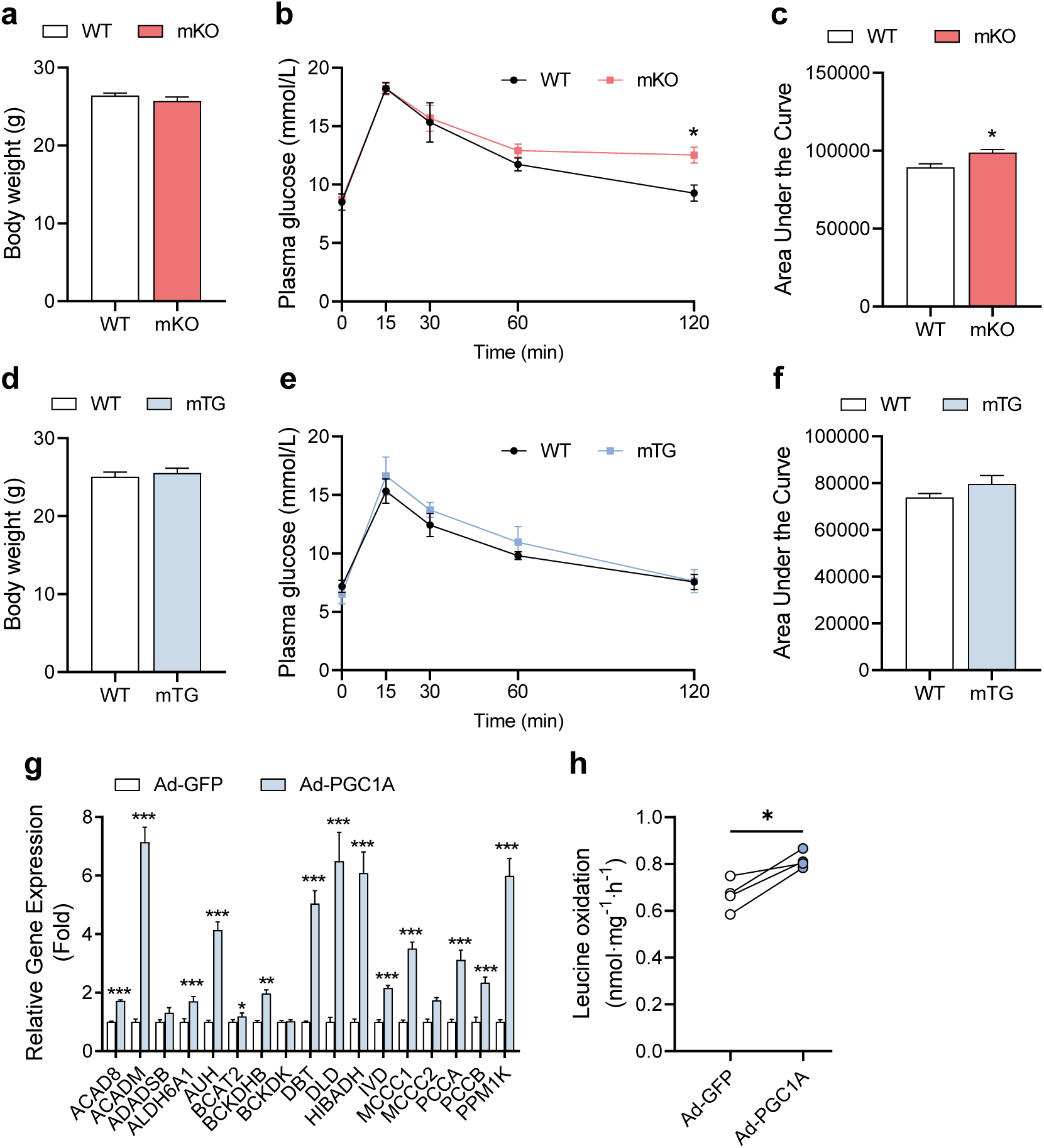
Mouse glucose tolerance and BCAA gene expression in *PPARGC1A* overexpressing C2C12 myotubes. **a)** Body weight of mKO mice and WT littermates (n=9). **b-c)** Oral glucose tolerance test and corresponding area under the curve of mKO animals and WT littermates (n=4). **d)** Body weight of mTG mice and WT littermates (n=6). **e-f)** Oral glucose tolerance test and corresponding area under the curve of mTG mice and WT littermates (n=4). **g)** BCAA gene expression in Ad-PGC1A C2C12 myotubes (n=4). **h)** Leucine oxidation in Ad-PGC1A C2C12 myotubes (n=4). *, *P*<0.05; **, *P*<0.01 and \*\*\**P*<0.001. Results are expressed as mean ± SEM. Ad-GFP, adenoviral overexpression of green fluorescent protein; Ad-PGC1A, adenoviral overexpression of PPARGC1A.

**ESM Figure 4 (Supplementary to Figure 6).**
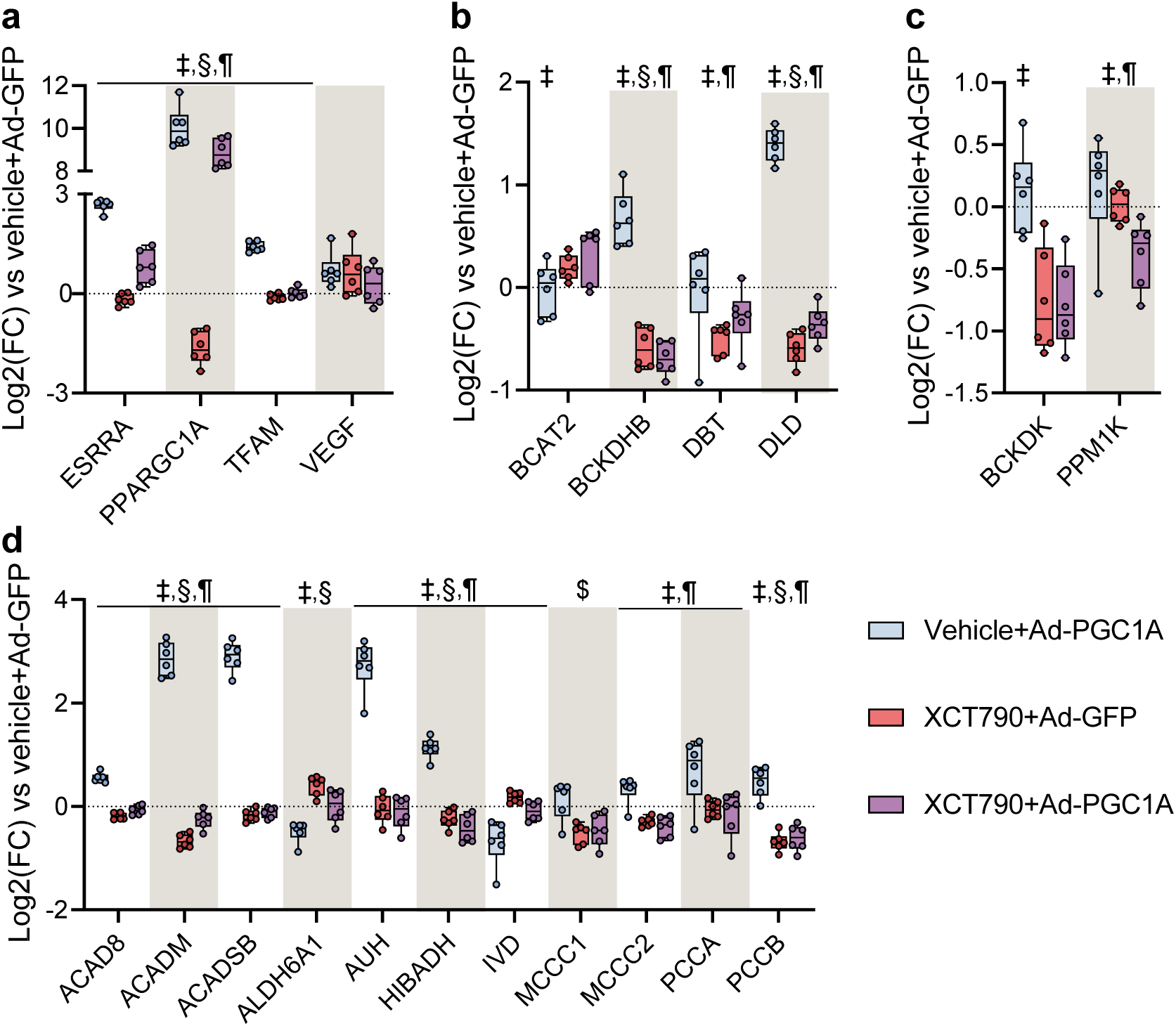
Effects of ERRα inverse agonist on BCAA gene expression. **a)** Expression of *ESRRA, PPARGC1A* and target genes in Ad-PGC1A cells treated with the ERRα inverse agonist XCT-790. **b-d)** BCAA gene expression in Ad-PGC1A cells treated with vehicle or XCT-790. Gene expression is shown as log2(fold-change) normalized to the scr+Ad-GFP (dotted line). Statistical analysis was performed using paired t-test (A-C, n=7) or two-way repeated measures ANOVA followed by Tukey’s *post-hoc* test (C-F, n=6). ‡, XCT-790 effect; §, Ad-PGC1A effect; ¶, interaction effect. siRNA, small interfering RNA; scr, scrambled siRNA; Ad-GFP, adenoviral overexpression of green fluorescent protein, Ad-PGC1A, adenoviral overexpression of *PPARGC1A*.

